# Consistency of superb microvascular imaging and contrast enhanced ultrasonography in detecting intraplaque neovascularization: a meta-analysis

**DOI:** 10.1101/2020.03.13.990374

**Authors:** Fang Yang, Cong Wang

**Author notes:** Corresponding author:Cong Wang, master,18098877102, Address: No.222 Zhongshan Road, Xigang District, Dalian City, Liaoning Province, China.

## Abstract

This meta-analysis aimed to identify the consistency of superb microvascular imaging(SMI) and contrast-enhanced ultrasonography(CEUS) in detecting intraplaque neovascularization(IPN). We searched PubMed, Web of Science, Cochrane Library, CISCOM, and CBM databases without language restrictions. Meta-analysis was conducted using STATA version 15.1 software. We calculated the pooled Kappa index. Ten studies that met all inclusion criteria were included in this meta-analysis. A total of 608 carotid plaques were assessed through both SMI and CEUS. The pooled summary Kappa index was 0.743(95 % CI=0.696-0.790) with statistical significance(z= 31.14, p<0.01). We found no evidence for publication bias (t=-1.21, p=0.261). Our meta-analysis indicates that SMI and CEUS display a good consistency in detecting IPN of carotid plaque, that is to say SMI ultrasound maybe a promising alternative to CEUS for detecting IPN of carotid plaque.

## Introduction

In today’s society, atherosclerosis incidence rate is high, and tends to be younger[1]. Atherosclerotic plaque can lead to carotid artery stenosis, affect the blood supply of carotid artery to the brain, and vulnerable carotid atherosclerotic plaques are prone to rupture and bleeding to form thrombus, which can enter the blood vessels of the brain with the blood, causing ischemic stroke events[2]. Stroke is a common refractory disease that seriously endangers human health and life safety[3]. The development of atherosclerotic plaque seriously affects the outcome and prognosis of the disease, and there is a significant consistency between IPN and atherosclerotic plaque vulnerability, so the IPN can be used as a high-risk factor to evaluate the vulnerability of plaque[4]. CEUS can visualize IPN effectively, but is an invasive examination requiring injection of contrast medium[5]. SMI is a new ultrasonic diagnosis technology which uses adaptive principle to display low-speed blood flow signal[6].Several studies had suggested that SMI, as a promising noninvasive alternative, can detect IPN with accuracy comparable to CEUS[7]. However, the results of these studies have been contradictory and the sample sizes were small. Therefore, we performed the present meta-analysis to identify the consistency of SMI and CEUS in detecting intraplaque IPN.

## Methods

### Literature search

We searched PubMed, Web of Science, Cochrane Library, CISCOM, and CBM databases without language restrictions. The following keywords and MeSH terms were used: [carotid] and [plaques or plaque or fatty Streak or fibroatheroma] and [contrast-enhanced ultrasound or contrast enhanced ultrasonography or contrast ultrasonography or ultrasound contrast imaging or CEUS] and [vulnerability or stability or neovascularization] and [superb microvascular imaging]. We also performed a manual search to find other potential articles.

### Selection criteria

The included studies must meet all four of the following criteria: (1) the study design must be a clinical cohort study, (2) the study must relate to the comparison of CEUS and SMI for detecting IPN, (3) intraplaque microvascular flow (IMVF) were be graded, and (4) published data in the row x column tables must be sufficient for Kappa index and standard error. If the study could not meet the inclusion criteria, it would be excluded. The most recent or the largest sample size publication was included when the authors published several studies using the same subjects.

### Data extraction

Relevant data were systematically extracted from all included studies by two researchers using a standardized form. There searchers collected the following data: year of article, the first author’s surname, sample size, number of IMVF grades, Kappa index, standard error, etc.

### Quality assessment

Methodological quality was independently assessed by two researchers according to a tool for the quality assessment of methodological index for non-randomized studies(MINORS). The MINORS criteria included 12 assessment items. Each of these items was scored as “yes” (2), “no” (0), or “unclear”(1). MINORS score ranged from 0 to 24; and score≥17 indicate a good quality.

### Statistical analysis

The STATA version 15.1 software(Stata Corporation, College Station, TX, USA) was used for Meta-analysis. We calculated the pooled summary Kappa index and its 95% confidence interval(CI). The Cochran’s Q-statistic and I^2^ test were used to evaluate potential heterogeneity between studies[8].If Q test shows a P<0.05 or I^2^ test exhibits>50% which indicates significant heterogeneity, the random-effect model was conducted, or else the fixed-effects model was used. In order to evaluate the influence of single study on the overall estimate, sensitivity analysis was performed. We conducted Begger’s funnel plots and Egger’s linear regression test to investigate publication bias[9].

## Results

### Characteristics of included studies

Initially, the searched keywords identified 65 articles. We Reviewed the titles and abstracts of all articles and excluded 42 articles; full texts and data integrity were also reviewed and 13 were further excluded. Finally, 10 studies that met all inclusion criteria were included in this meta-analysis [10-19]. Figure1 showed the selection process of eligible articles. A total of 608 carotid plaques were assessed through both SMI and CEUS. MINORS scores of all included studies were 17. We summarized the study characteristics and methodological quality in Table1.

**Fig. 1.**
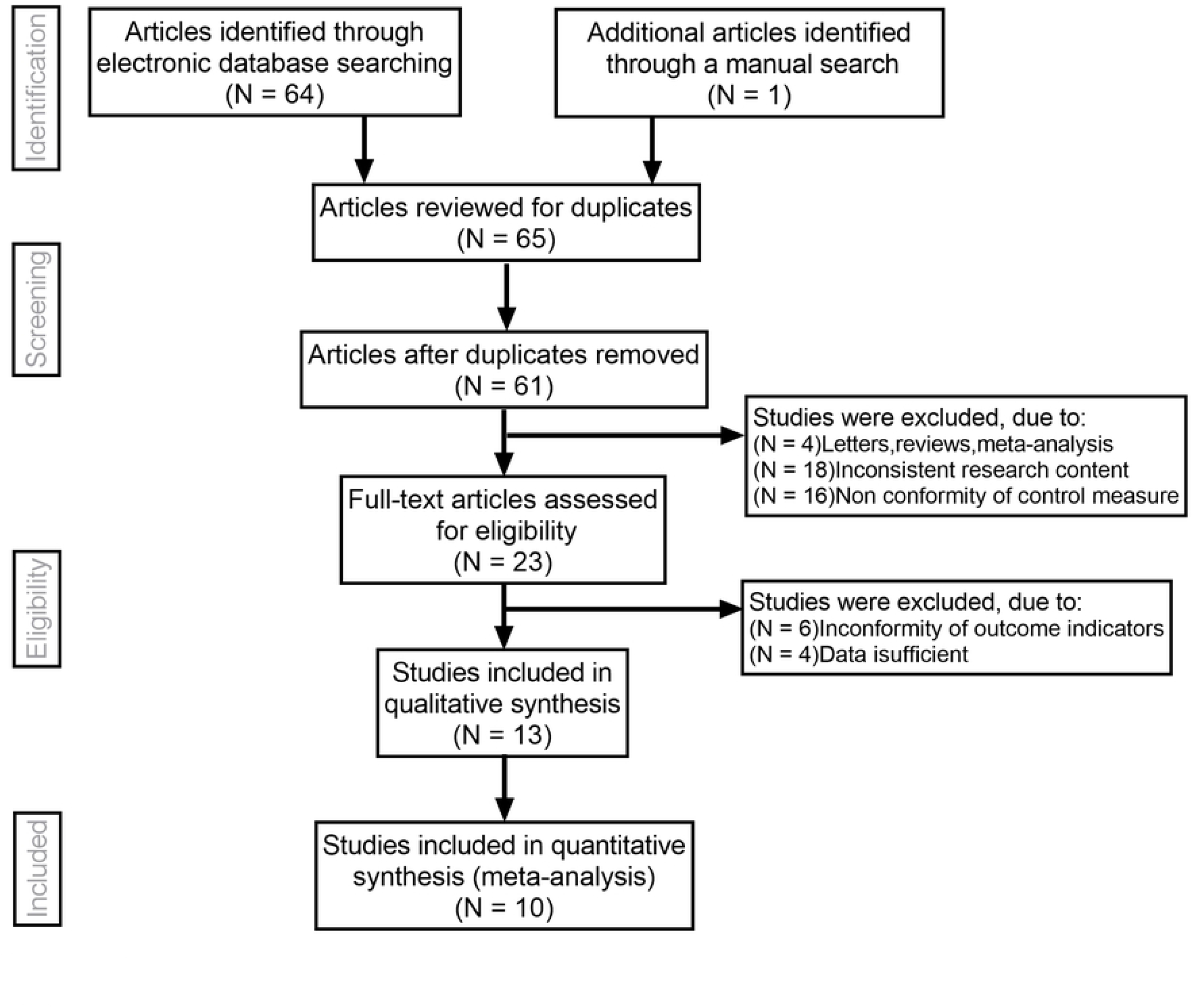
Flow chart of literature search and study selection. Ten studies were included in this meta-analysis.

### Quantitative data synthesis

The fixed effects model was used due to not obvious heterogeneity among the studies(I^2^=0%,P=0.531). Sensitivity analysis was carried out, and none of them caused obvious interference to the results of this meta-analysis(Figure2). The pooled summary Kappa index was 0.743(95 % CI=0.696-0.790) with statistical significance(z= 31.14, p<0.01), which indicated that SMI and CEUS have a good consistency in detecting IPN of carotid plaque(Figure3). We found no evidence of obvious asymmetry in the Begger’s funnel plots(Figure4). Egger’s test also did not display strong statistical evidence for publication bias(t=1.21, p=0.261).

**Fig. 2.**
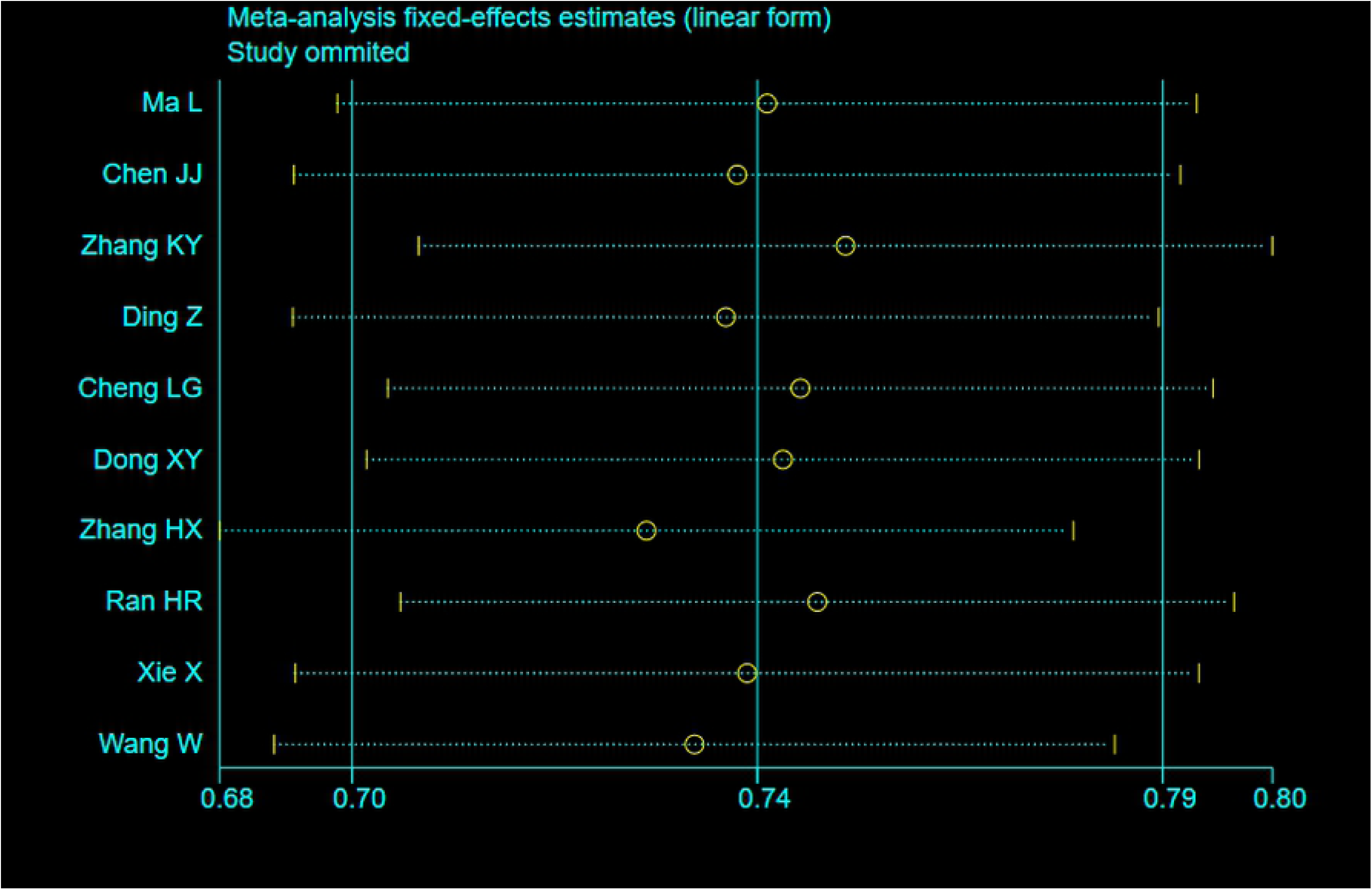
Sensitivity analysis. None of them caused obvious interference to the results.

**Fig. 3.**
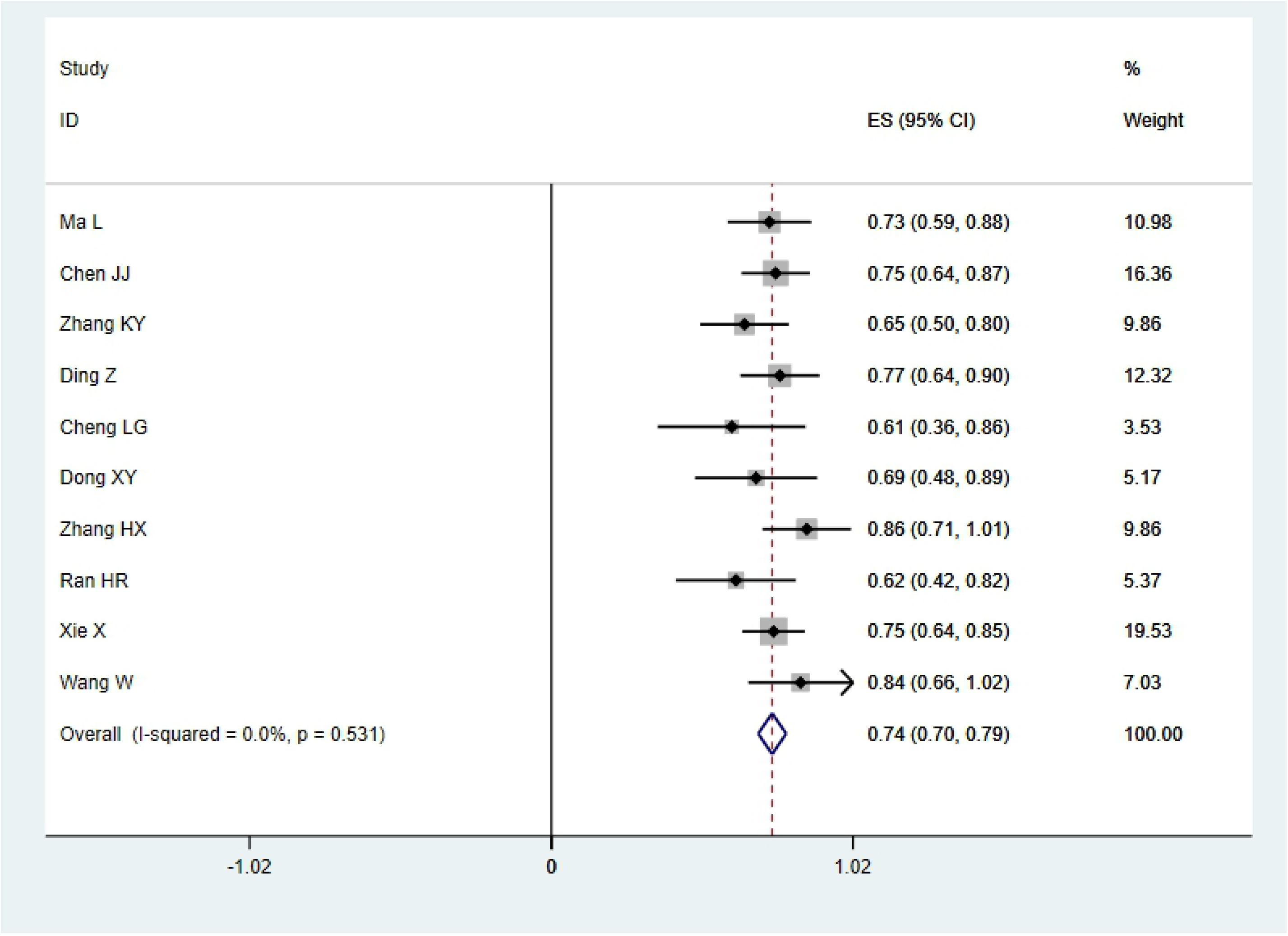
Forest plots of Kappa index for SMI in the detection of IPN comparable to CEUS.

**Fig. 4.**
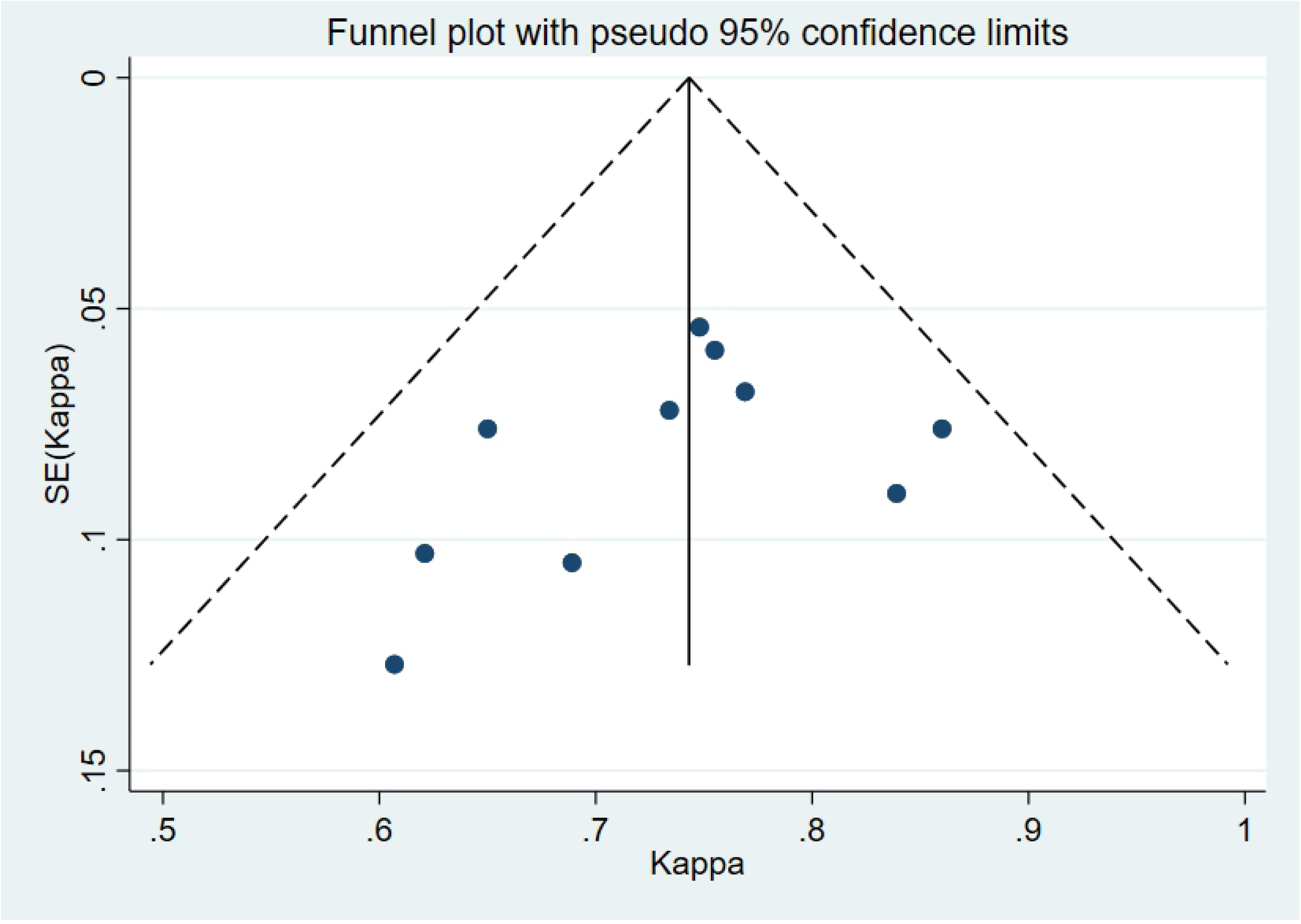
Begger’s funnel plot of publication bias on the pooled Kappa. No publication bias was detected in this meta-analysis.

## Discussion

Carotid atherosclerotic plaque is closely related to cerebrovascular events, and its direct factor is the structure and composition of plaque, which indicates the occurrence and development of ischemic cerebrovascular events in the future[20]. Those unstable plaques that are easy to rupture, fall off and cause distal embolism are called vulnerable plaques[21]. The morphological characteristics of vulnerable plaque include the following characteristics: irregular plaque surface or ulcer formation, thin fiber cap or fissure, bleeding, rich lipid and inflammatory active ingredients, and neovascularization in plaque[22]. Neovascularization plays a central role in plaque initiation, progression and rupture, and is a predictor of plaque instability and stroke risk[23]. Histopathological evidence shows that compared with relatively stable plaque, the presence and density distribution of neovascularization in plaque are closely related to plaque rupture, and neovascularization is often located in the fibrous cap fissure area, lipid enrichment area, inflammatory active area. The detection of neovascularization in plaque can effectively evaluate the stability of atherosclerotic plaque and even predict the occurrence of cardiovascular and cerebrovascular diseases[24]. Finding a convenient, safe and reproducible imaging method to detect the stability of arterial plaque has always been the focus of clinical research.

Conventional two-dimensional ultrasound and color Doppler ultrasound can observe and measure the echo, shape and thickness of plaque, and evaluate the degree of vascular stenosis caused by plaque, but they can not judge and evaluate the stability of plaque comprehensively and accurately, while CEUS has high spatial and temporal resolution, and microbubbles have the same fluidity as red blood cells, and some scholars use CEUS to detect neovascularization in plaque as reliable evidence for the diagnosis of vulnerable plaque[25, 26]. However, because of the high cost of contrast media, trauma examination and the risk of allergy of contrast media, CEUS is limited to some extent. Therefore, it is necessary to explore a simple, noninvasive and inexpensive ultrasound examination method.

SMI technology is based on the high-resolution Doppler technology, using the up-market ultrasonic diagnostic equipment of Aplio series to build “high-density beamformer” and “real-time application platform”, and imaging the low flow velocity blood flow with a higher frame rate, while traditional Doppler ultrasound uses filtering technology to eliminate noise and motion artifacts, resulting in the loss of low-speed blood flow information. SMI technology can identify the noise generated by blood flow and tissue movement, and use adaptive calculation method to display the real blood flow information, so that the low-speed blood flow signals can be eparated and displayed from filtered clutter signals[27]. Present studies have revealed that SMI, as a simpler, safer, cheaper, and noninvasive technique may facilitate the visualization of carotid artery IPN without the use of a contrast agent[7, 14]. However, quantitative evaluations of IMVF signal in SMI have not been established and the relationship between SMI findings and degree of enhancement on CEUS remains unclear. At present, there is a lack of multi center and large sample research in this aspect. This study aims to provide a comprehensive and reliable conclusion on the consistency of SMI and CEUS in detecting IPN.

In the present meta-analysis, we systematically evaluated the technical performance and the consistency of SMI and CEUS in detecting IPN. Finally, 10 independent studies were included with a total of 608 carotid plaques assessed. The pooled summary Kappa index was 0.743 with statistical significance. Furthermore, our results found no direct evidence for publication bias. Taken together, consistent with previous studies, our findings strongly suggest that SMI and CEUS display a good consistency in detecting IPN of carotid plaque, that is to say SMI ultrasound maybe a promising alternative to CEUS for detecting IPN of carotid plaque.

However, this meta-analysis still has some limitations. Firstly, our results had lacked sufficient statistical power due to relatively small sample size and low-quality included studies. On the other hand, meta-analysis is a retrospective study that may lead to subject selection bias. Thirdly, this meta-analysis failed to obtain original data from the included studies, which may limit further clinical assessment of values of SMI in detecting IPN. Importantly, the majority of included studies originated from China, which may adversely affect the reliability and validity of our results.

In conclusion, our meta-analysis suggests that SMI and CEUS display a good consistency in detecting IPN. However, due to the limitations mentioned above, further detailed studies are still required to confirm our findings.

## Ethical statements

All procedures followed were in accordance with the ethical standards of the responsible committee on human experimentation (institutional and national) and with the Helsinki Declaration of 1964 and later versions.

Informed consent was obtained from all patients for being included in the study.

## Acknowledgments

We would like to acknowledge the helpful comments on this paper received from our reviewers. Also, we would like to thank all our colleagues working in the Ultrasound Department of The First Affiliated Hospital to Dalian Medical University.

